# *DeepStomata:* Facial Recognition Technology for Automated Stomatal Aperture Measurement

**DOI:** 10.1101/365098

**Authors:** Yosuke Toda, Shigeo Toh, Gildas Bourdais, Silke Robatzek, Dan Maclean, Toshinori Kinoshita

## Abstract

Stomata are an attractive model for studying the physiological responses of plants to various environmental stimuli^1–3^. Of the morphological parameters that represent the degree of stomatal opening, the length of the minor axis of the stomatal pore (the stomatal aperture) has been most commonly used to dissect the molecular basis of its regulation. Measuring stomatal apertures is time consuming and labour intensive, preventing their use in large-scale studies. Here, we completely automated this process by developing a program called *DeepStomata*, which combines stomatal region detection and pore isolation by image segmentation. The former, which comprises histograms of oriented gradients (HOG)-based stomatal detection and the convolutional neural network (CNN)-based classification of open/closed-state stomata, acts as an efficient conditional branch in the workflow to selectively quantify the pores of open stomata. An analysis of batches of images showed that the accuracy of our automated aperture measurements was equivalent to that of manual measurements, however had higher sensitivity (i,e., lower false negative rate) and the process speed was at least 80 times faster. The outstanding performance of our proposed method for automating a laborious and repetitive task will allow researchers to focus on deciphering complex phenomena.

---

To fully understand the molecular mechanisms underlying the regulation of stomatal movement, sufficient numbers of stomata must be analysed under a range of experimental conditions. To date, ocular micrometer-assisted visual estimation has been widely used to measure stomatal apertures; however, as stomatal movements are highly sensitive to their environment, the number of stomata that can be analysed before their apertures change is very limited using this method. Imaging intact or replica leaf surfaces enables the analysis to be deferred, alleviating the possibility of aperture changes during observation; however, aperture measurements are still labour intensive. Recent technical advances in computer vision can assist the quantification process. CARTA, an ImageJ plugin that utilizes a self-organizing map algorithm, allowed the semi-automatic classification of open/closed-state stomata^4^. More recently, a script that automatically quantifies stomatal apertures has been developed (https://github.com/TeamMacLean/stomatameasurer); however, it requires that images be taken using a confocal scanning laser microscope with a high-contrast background. In many studies, images are acquired using an optical camera with bright-field microscopy, which contains dense and complex pixel information. To the best of our knowledge, no methods currently facilitate the automatic measurement of stomatal apertures from such images.

As a solution to these issues, phenotypes that can serve as a proxy for stomatal opening have been used to identify factors that regulate stomatal movement. Thermal imaging of leaves or monitoring their water loss can reflect the stomatal aperture-dependent transpiration rate^5,6^, while immunostaining can detect the degree of H^+^-ATPase phosphorylation that provides the motive force for stomatal opening^7,8^. These approaches have been successfully used to isolate genetic factors or compounds involved in stomatal regulation; however, these proxies do not always reflect the true status of the stomatal aperture. Recently, we isolated compounds that regulate stomatal movements (stomatal closing compounds; SCLs) in a chemical screen focussed on stomatal apertures^9^. Limiting to include only chemicals for quantification that altered stomatal aperture greatly enhanced the screening throughput to 150 compounds per day, which resulted in the successful identification of 9 candidates from over 20,000 chemicals. However, we could not exclude the possibility of overlooking false negatives due to technical error or a gradual effect of the compound. Automating the quantification process would not only ensure high throughput, but would also provide uniform quality and higher sensitivity in various analyses. Here, we propose a complete automation of this process, enabling the batch processing of bright-field images containing both open and closed stomata (Fig. 1).

**Figure 1.**
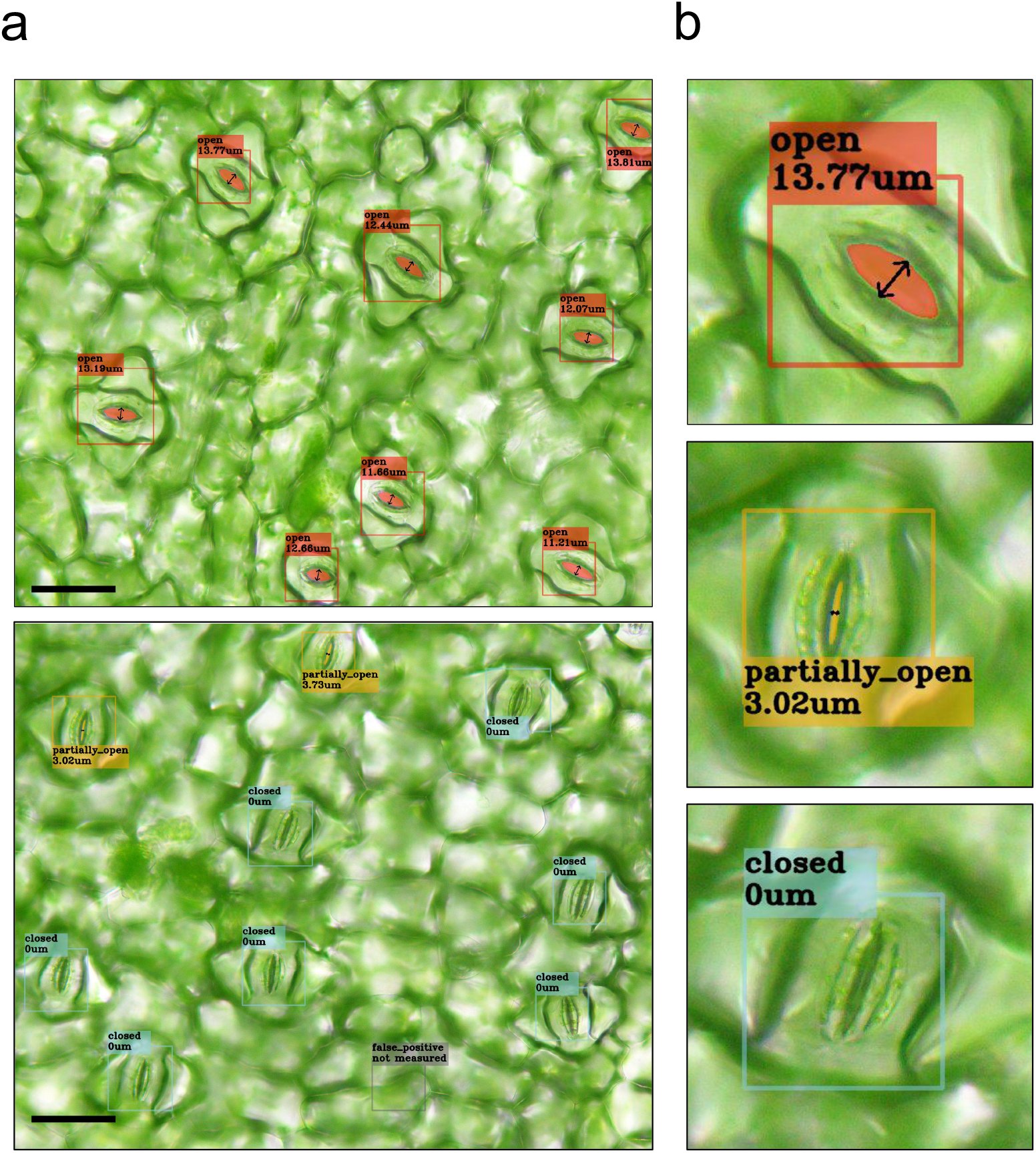
Automatic quantification of stomatal aperture by *DeepStomata*. **a**, Taking bright-field images as an input, our proposed method enables the automation of stomatal aperture measurement. The detected stomata and their open, partially open, or closed states are marked as red, orange, and blue boxes, respectively. Enlarged views of representative stomata from each image are shown in **b**. For each stoma, its open, partially open, or closed status, and its quantified aperture are shown. The detected stomatal pore regions of the open stomata are coloured red or orange.

During conceptualization, we were inspired by the principle of facial recognition^10,11^ (Fig. 2A), or more broadly region proposal algorithms. First, the position of the face within the image is isolated either using a predefined threshold, or by a machine learning-based method such as histograms of oriented gradients (HOG), local binary patterns, or Haar-like cascades. Next, using the cropped face image as an input, high-context information such as age, biological sex, emotion, and identity are further qualitatively or quantitatively evaluated using a convolutional neural network (CNN). CNNs are a type of neural network used in deep learning, and are powerful tools for image recognition and classification^12^. Since deep learning enables the neural network itself to learn the most suitable feature, it generally prevails over conventional machine learning in various situations. In this report, we propose a method for stomatal aperture measurement that partially follows facial recognition: (1) Detection of the stomatal candidate regions within the image using a HOG; (2) Classification of whether the detected stomata are open or closed using the CNN; and (3) Selective quantification of the apertures of open stomata by pore region isolation using binary based image segmentation.

**Figure 2.**
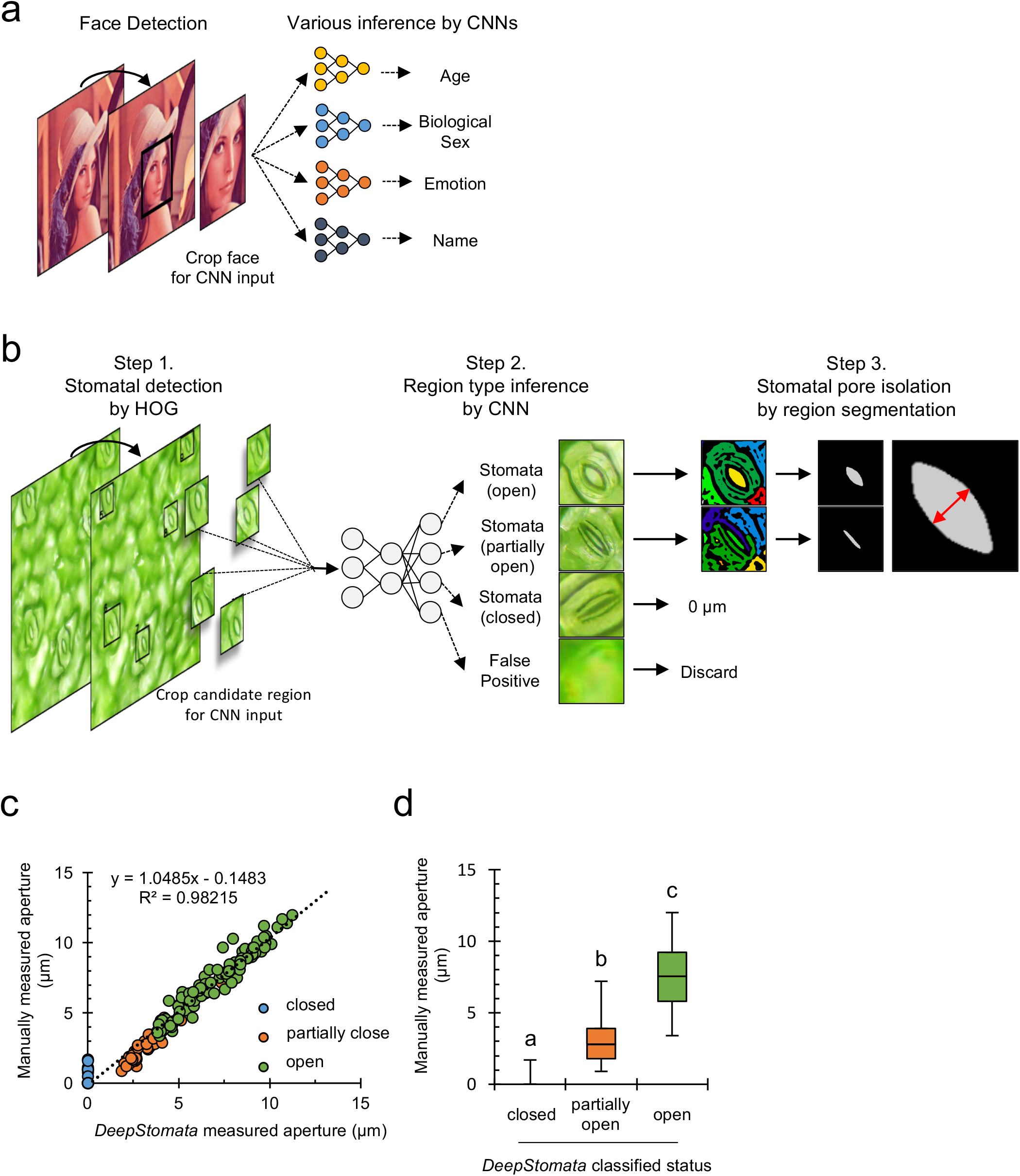
Workflow and processing accuracy of automatic stomatal aperture quantification. **a**, Schematic diagram of facial detection. First, the sub-region of an image that contains a face is detected and cropped (Face Detection) using predefined features or machine learning. Next, the cropped images are passed to the trained convolutional neural networks (CNNs) to infer complex factors such as age, biological sex, emotions, or identity (name). **b**, Schematic diagram of our proposed method. First, the subregions that contain stomata are obtained by the HOG detector (Step 1; See Supplementary Figure 1a for details of the constructed HOG detector). Next, cropped images are processed by the CNN to infer whether the respective input image is an open, partially open, or closed stoma, or a false positive that does not contain a stoma (Step 2; See Supplementary Figure 2a for details of the CNN architecture). Images classified as false positives are discarded from the analysis. Closed stomata are assigned a 0-µm stomatal aperture. Open and partially open stomata are passed to the pore quantification step (Step 3), in which the images are processed to create and label their segmented regions. To discriminate the stomatal pore, manually set criteria were introduced to filter out non-pore regions (See Methods for details). **c**, Scatter plot of automatically quantified stomatal apertures versus manually quantified apertures. The classifications denoted by the CNN are displayed in different colours. The equation and R-squared value of the regression line are displayed. Data categorized by classification are summarized in **d**. **d**, Box plots showing the stomatal apertures of the closed, partially open, and open stomata classified by the CNN. Statistical differences were determined using a one-way ANOVA followed by a Tukey post-hoc analysis; different letters indicate significant differences (p < 0.01).

The pipeline of the proposed method is summarized in Fig. 2b. The details of the respective processes are provided in the Methods section of this paper. For stomatal detection, we combined HOG with a max-margin object detection method^13,14^. The constructed HOG detector scans the input image and the HOGs are compared within the sub-region of images for stomatal detection (Supplementary Fig. 1a). We constructed a detector that can accept any orientation and degree of stomatal aperture (Supplementary Fig. 1b). The precision and recall value of the detector on the test dataset were 0.954 and 0.940, respectively (Supplementary Fig. 1c). In addition to stomata, the detector was found to collect false positives, images containing a background non-stomata region; therefore, we categorized the collected images into four classes: open, partially open, or closed stomata, or false positives. We then trained a CNN that takes these images as an input and infers the probability of each belonging to the four separate classes (Supplementary Fig. 2a). After training, the classification accuracy reached an average of 92.9% on the test dataset (Supplementary Fig. 2b). Images classified as false positives were discarded, and closed stomata were assigned a stomatal aperture of 0 µm. Regions containing open and partially open stomata were further analysed to isolate the stomatal pore. This was done by combining region segmentation with the filtering out of non-pore regions using manually defined geometrical shape parameters. Finally, the minor axis length of the stomatal pore was retrieved as the stomatal aperture.

To evaluate the accuracy of the proposed method, the automatically measured stomatal apertures were compared with manually measured values. The slope (1.048) and the R-squared value (0.982) of the regression line revealed a negligible difference between the two datasets (Fig. 2c). In addition, the status of the stomata inferred by the CNN showed a clear segregation between open, partially open, and closed stomata, based on their stomatal apertures. The maximum aperture of the closed stomata was 1.7 µm. The minimum and maximum values of the partially open stomata were 0.9 µm and 3.4 µm, respectively, while for the open stomata they were 7.2 µm and 12 µm, respectively (Fig. 2d). This suggests that deep learning not only functions as an efficient conditional branch, but also enables a human-like qualitative decision to be embedded in the image analysis pipeline.

Based on the proposed method, we built a program (*DeepStomata*) that enables batch image processing. We first analysed the effect of the plant hormone abscisic acid (ABA) and the fungal toxin fusicoccin (FC) on stomatal movement, which are known to inhibit and induce stomatal opening, respectively^1,2,15^. We determined their half-maximal active concentration by treating stomata with various concentrations of the compounds, followed by image acquisition and automatic quantification (Fig. 3a and 3b). The calculated half-maximal inhibitory concentration (IC_50_) of ABA and the half-maximal effective concentration (EC_50_) of FC against stomatal opening were 474 nM and 4.46 µM, respectively. Our calculated IC_50_ of ABA was close to a previously reported value, 0.36 µM^9^. Next, we reanalysed 715 images acquired from our recent chemical screen to identify compounds that alter stomatal movement^9^. In addition to the stomatal apertures, we also calculated the percentage of open, partially open, and closed stomata classified by the CNN. Of the 82 compounds evaluated (Fig. 3c,e and Supplementary Fig. 3), two (SCL1 and SCL1-2) inhibited light-induced stomatal opening (mean stomatal aperture below 50% of DMSO-treated plants; Student’s *t*-test p < 0.05) (Fig. 3d, dark blue). We further confirmed that the stomatal apertures measured by our program were identical to those measured manually following treatment with these compounds (Fig. 3e). SCL1 and SCL1-2 were the only bioactive compounds identified in the previous study^9^; however, with a more marginal threshold (mean stomatal aperture below 80% of the DMSO-treated plants; Student’s *t*-test p < 0.05), eight other compounds were also found to have an inhibitory effect on stomatal opening (Fig. 3d, light blue). These compounds were overlooked in the qualitative evaluation used in our previous screen. Moreover, the total time required to process the dataset was 12.3 minutes in the present study, while it was estimated to take over 16 hours for a single worker to manually measure the apertures in the 715 images at a constant speed without any breaks (Fig. 3f). Collectively, these results suggest that introducing automation by *DeepStomata* enables a higher sensitivity (i.e., a lower false negative rate) and a dramatically faster analysis.

**Figure 3.**
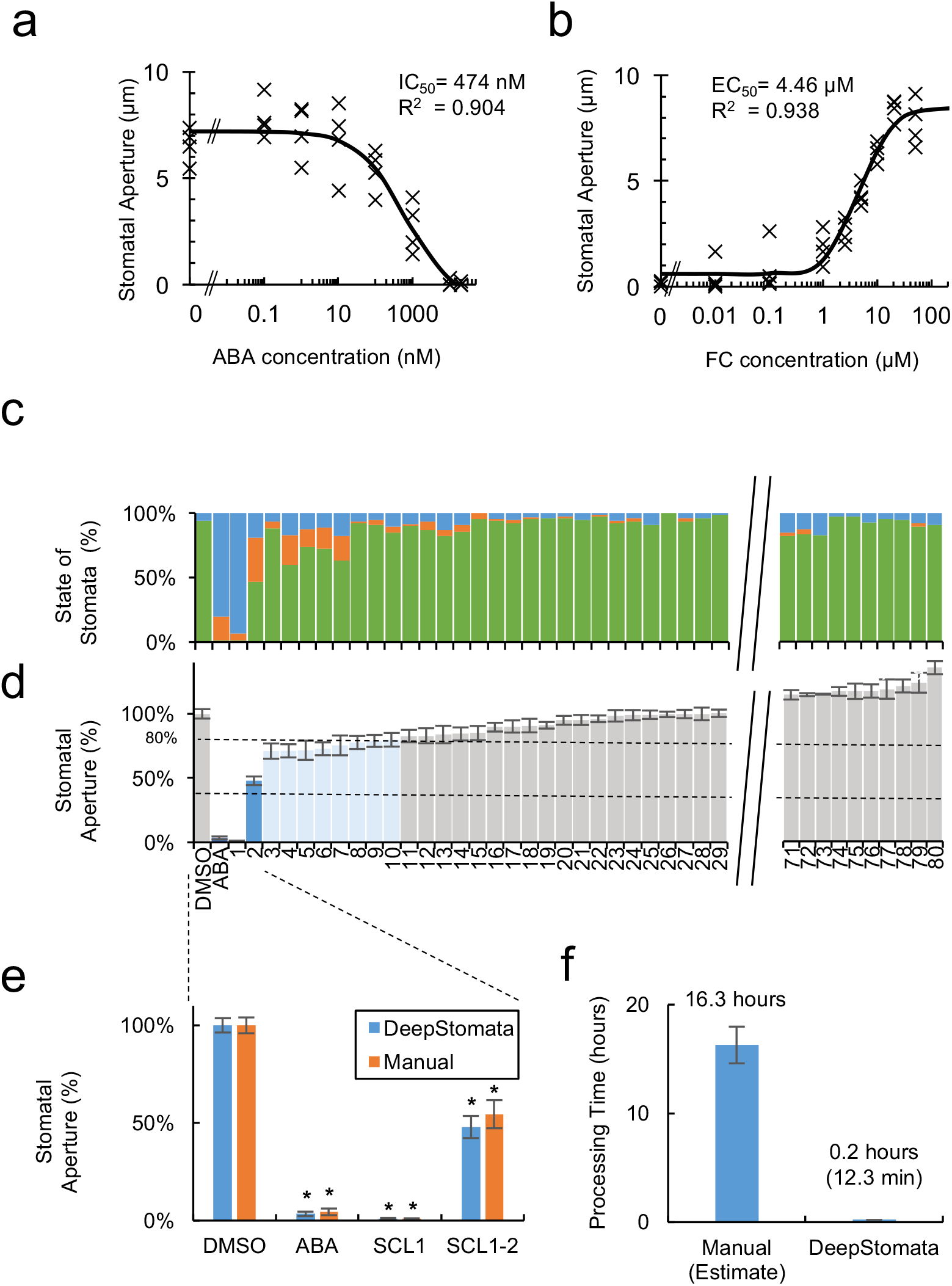
Examples of applications in the characterization and screening of compounds that control stomatal movements. **a–b**, Dose-response curves of the inhibition of light-induced stomatal opening by abscisic acid (ABA) (**a**) or promotion of stomatal opening in the dark by fusicoccin (FC) (**b**). The IC_50_ or EC_50_ value and the R-squared value of the sigmoid curves are displayed. **c–e**, Summary of the chemical screen performed to identify compounds that inhibit light-induced stomatal opening, analysed using *DeepStomata*. Dayflower leaf discs were treated with either DMSO, 10 µM ABA, or 50 µM of a compound from the chemical library, and then irradiated under red light and blue light for four hours^9^. For clarity, some of the results have been omitted. Refer to Supplementary Fig. 3 for the complete set of results. **c**, Stacked bar graph representing the ratio of open (green), partially open (blue), and closed (orange) stomata classified by the CNN. **d–e**, Values of stomatal apertures relative to those following DMSO treatment. Asterisks indicate significant differences to the DMSO treatment (Student’s *t*-test, p < 0.05). Two compounds that caused more than a 50% inhibition of stomatal opening (SCL1 and SCL1-2; p < 0.05) are coloured dark blue, while eight compounds that caused more than a 20% inhibition of stomatal opening (p < 0.05) are coloured light blue. **e**, Manually and automatically measured stomatal apertures of plants treated with SCL1 and SCL1-2. Means ± SE (n ≥ 60 stomata). **f**, Comparison of the time required to quantify the 715 images in this analysis using either *DeepStomata* (Automation) or manual quantification (Manual). For the manual quantification, the time was estimated from the average time taken to analyse 10 images multiplied by the number of images in the dataset.

As reviewed by Grys et al.^16^, the phenotypic analysis of biological images first requires that the object of interest be isolated from the background. Thresholding or edge-detection algorithms followed by region segmentation have often been used to isolate cells or cellular components; however, the input previously had to be free of background noise, such as an image acquired using a confocal laser scanning microscope. Such approaches were not suitable for evaluating stomatal pores in our images, which depict complex leaf surfaces and contain rich information in the non-stomatal regions (Fig. 1). Moreover, the lack of apparent pores in the closed stomata made their automated quantification difficult. In this study, we utilized HOG-based stomata detection with a CNN-based object classifier, which was inspired by a region proposal method. This allowed us not only to count the closed stomata, but also to crop the sub-regions containing open stomata, making the pore isolation more straightforward using region segmentation.

In the present study, we established a method that enables the complete automation of stomatal aperture measurement. To the best of our knowledge, this is the first approach that can process stomata images obtained using a bright-field ocular microscope. Our study also demonstrates that techniques used in the field of computer vision and pattern recognition can have powerful applications in plant image analysis. Moreover, recent technical advances in deep learning have enabled the direct detection of object positions^17–20^ and segment regions at a pixel level^21,22^, in addition to their classification. Since such methods are capable of learning and processing more complex tasks than conventional machine learning approaches, we expect that programs incorporating these techniques will further broaden the versatility of automated tasks used for laborious and repetitive plant image analyses, such as chemical/genetic screening, thus freeing researchers to focus on deciphering complex phenomena.

## Methods

### Code Availability

*DeepStomata* was written in Python with open source libraries. The source code is available in github repository (https://github.com/totti0223/deepstomata/) under MIT license.

### Stomatal Detection by HOG

Stomatal detection by HOG was performed using the *simple_object_detector* module of the dlib library (http://dlib.net), based on the developer’s tutorial (http://dlib.net/train_object_detector.py.html). Greyscaled images were used for training and detection. Notably, upon detector training and detection, the input image was down-sampled (2448×1920 px to 512×402 px) to eliminate some of the minor noise within the original image, which could affect the detection accuracy, and to enhance the detection speed. The coordinates of the detected objects in the down-sampled image were then converted for mapping to the original image by multiplying by the scaling factor, 4.78 (2448 px / 512 px). Sub-regions were cropped according to the co-ordinates and passed to the following classifier.

### Region Classification by CNN

A TensorFlow library (https://www.tensorflow.org/) was used to construct the CNN classifier, the architecture of which is described in Supplementary Fig. 2a. A dataset containing, respectively, 590, 480, and 799 images of open, partially open, and closed stomata and 99 false positives was prepared for training the network. Images were collected by manually classifying the images obtained by the HOG detector. Since the primary objective was to discriminate between open and closed stomata and exclude the false positives, no stringent criteria were introduced to discriminate between open and partially open stomata. The partially open class was defined to examine whether the CNN could learn a human-like classification. Stomata were defined as open if their stomatal apertures were larger than about 4 µm, and defined as partially open if they had smaller apertures.

The classified images were resized to 56×56 px to train the CNN. Training was performed by minimizing the loss value using the Adam optimizer algorithm with the following parameters: 6% drop out rate; 0.0001 learning rate; 5000 iterations; and 200 batch size. If the maximum probability value of the classifier showed that the image depicted an open or partially open stoma, the image was passed to the final quantification step. In the case of images classified as closed, the stomatal apertures of the objects were assigned a value of 0 µm. False positives were discarded.

### Stomatal Pore Quantification

The apertures of the stomatal pores were quantified using the image processing modules of scikit-image (http://scikit-image.org). The input image, the cropped image generated by the detector, was first converted to greyscale, Gaussian blurred, and binarized using an adaptive threshold. Erosion and dilation followed by dilation and erosion were applied to remove minor noise and fill gaps. Next, the *ndimage.label* module of scipy (https://www.scipy.org) and the *measure.regionprops* module of scikit-image were used to label and retrieve the geometric properties of the segmented regions. To discriminate the stomatal pore, manually defined criteria (area, solidity, and major axis length) were introduced to filter out non-stomatal pore regions. The x,y co-ordinates of the centroid of the respective regions were also introduced as filtering criteria because the pore tended to be located at the centre of the detected region. The optimized parameters for filtering were: 100 px < area < 6000 px, solidity > 0.8, major axis length < 40 px, and an x,y co-ordinate of 0.2 × (X,Y) < (x,y) < 0.8 ± (X,Y) for the centroid, where (X,Y) is the width and height of the cropped sub-region. The minor axis length of the remaining region was retrieved as the value of the stomatal aperture.

### Image Acquisition

Dayflower (*Commelina benghalensis*) plants were grown in soil in a glasshouse at 25°C, under a natural photoperiod of 10 h light/14 h dark. Dayflower leaf discs were treated with chemical compounds for screening as described previously^9^. Briefly, plants were incubated in darkness overnight to ensure the complete closure of their stomata. Using a biopsy punch (Kai Medical), 4-mm leaf discs were excised from fully expanded leaves under dim light. The discs were emerged in buffer (50 mM KCl, 0.1 mM CaCl_2_, 0.4 M mannitol, 5 mM MES-KOH, pH 6.5) with or without a screening compound and with or without light irradiation [150 µmol m^−2^ s^−1^ red light (LED-R; Eyela), 50 µmol m^−2^ s^−1^ blue light (Stick-B-32; Eyela), or white light (LED unit of BX43 microscope; Olympus)]. Images of the leaf disc were taken using a stereomicroscope (BX43; Olympus) with a 10× objective lens (UPlanFL N; Olympus) and a CCD camera (DP27; Olympus). Upon acquisition, the “Extended Focus Imaging” function of the cellSens standard software (Olympus) was used to maximize the number of fully focused stomata within each image. The stomatal apertures were manually quantified using cellSens standard software. Closed stomata were assigned an aperture of 0 µm, and the stomatal apertures of the open stomata were quantified by measuring the minor axis length of the stomatal pore.

## Acknowledgements

We would like to thank M. Uchida (Nagoya University) and E. Asai (Nagoya University) for their assistance in acquiring the images used in part of this study, and W. Ye (Nagoya University) for critically reading the manuscript. We thank A. Sato (Chemical Library Center of ITbM, Nagoya University) for providing information of chemical library compounds. We thank T. Suzuki (Chubu University) and the Global COE program “Advanced Systems-Biology” of Nagoya University for the intensive bioinformatics seminar. This research was supported by a Japan Science and Technology Agency (JST) PRESTO grant (Y.T., JPMJPR1705); the Advanced Low Carbon Technology Research and Development Program of JST; the Ministry of Education, Culture, Sports, Science and Technology of Japan [Scientific Research on Priority Areas (T.K., 15H059556; 15K21750)].

## Author contributions

Y.T. and T.K. directed the study and wrote the manuscript. Y.T. designed and wrote the program code with assistance from D.M. G.B., S.R., and D.M. conceptualized the image analysis. Y.T. and S.T. collected and directed the manual quantification of stomatal apertures from the microscopy images.

## Additional information

Supplementary information is available online. Correspondence and requests for materials should be addressed to Y.T. and T.K.

## Figure Legends

**Supplementary Figure 1.**
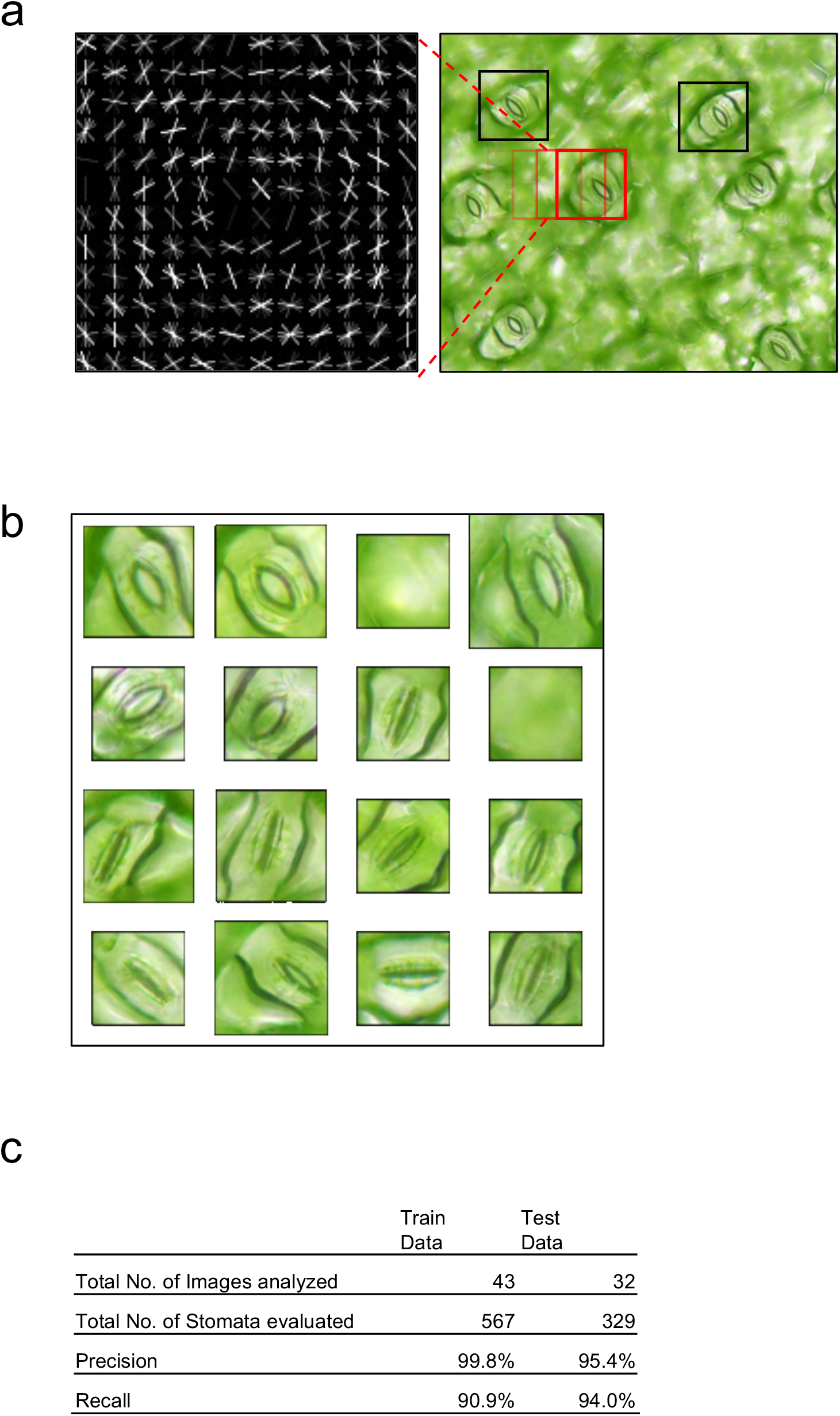
Details of the histogram of oriented gradient (HOG)-based stomatal detection. **a**, The trained HOG detector (left) was used to locate the coordinates of the stomata within the image (right) using a sliding window (red box). The detected stomata are marked with black boxes. **b**, Example of images retrieved by the HOG detector, including two regions that do not contain stomata (false positives). **c**, Precision and recall values of the HOG detector against the training and test data. Training data were used to construct the detector, while the test data were independently prepared to evaluate the detector’s accuracy.

**Supplementary Figure 2.**
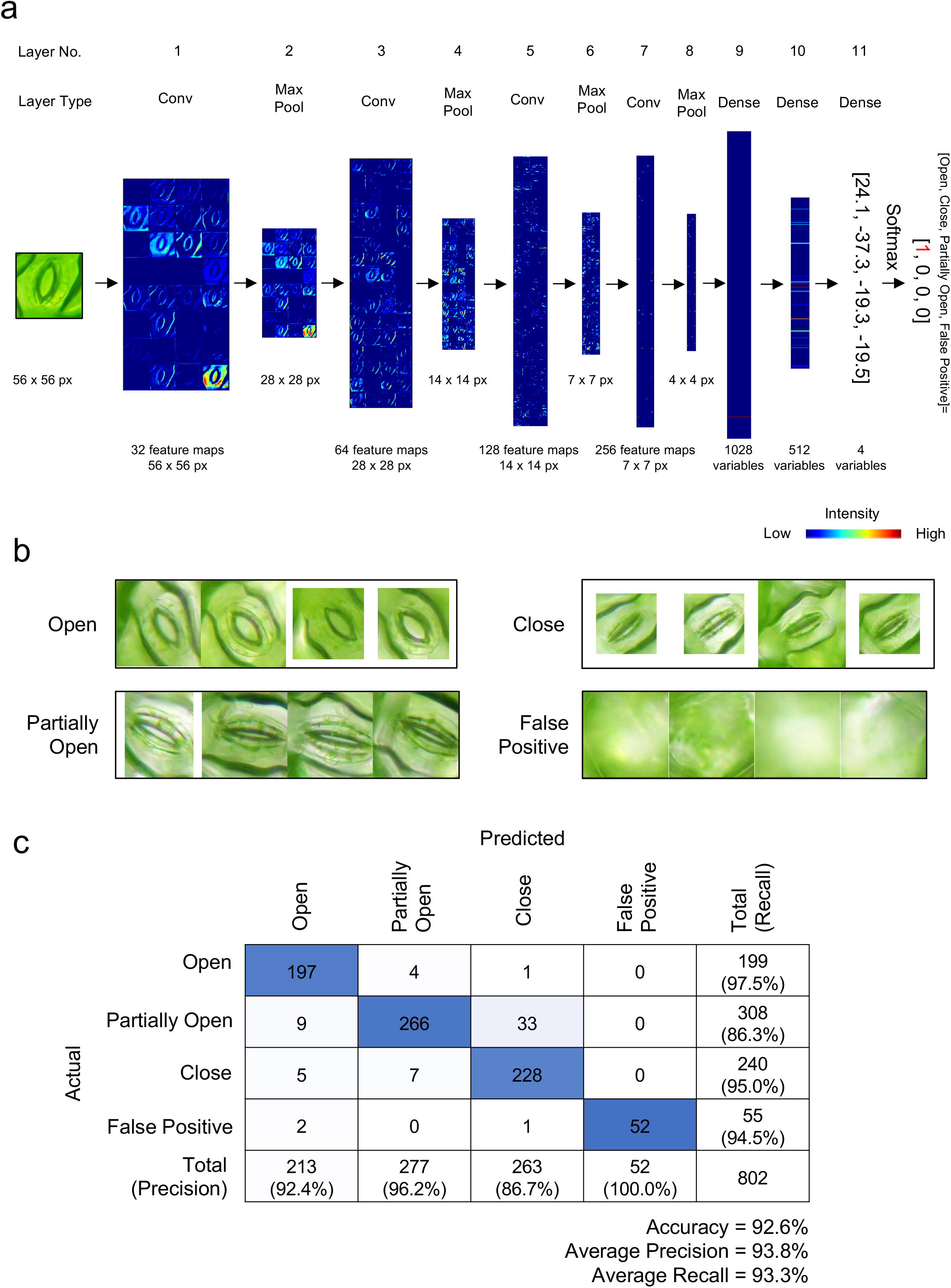
Details of convolutional neural network (CNN)-based object classification. **a**, Architecture of the convolutional neural network (CNN) classifier used in this study. Here, one image is processed through the CNN classifier, resulting in a four-vector variable in layer No. 11. The softmax function is next applied to truncate the sum of the values to 1, which represents the probability values that the image contains an open, closed, or partially open stoma, or a false positive. In this example, the input image is converted to [1,0,0,0], indicating that the input image is most probably an open stoma. In each of the convolutional layers, input images are convoluted using a weight with a kernel size of 3 × 3 px and a stride of 1 × 1 px, bias addition, followed by truncation using a rectified linear unit function. Max pooling was performed using a stride of 2 × 2 px and a kernel size of 3 × 3 px. The input image, intermediate images, and the variables produced within the CNN are displayed. Layer No. and Layer Type [Conv, convolutional layer; Max Pool, max pooling layer; Dense, fully connected layer] are displayed above each image. The pixel size of the input image, the intermediate data, and the number of weights or variables used in each layer are displayed below each image. **b**, Example of images classified using the CNN. **c**, Multi-class confusion matrix for the classification results obtained with CNN in the test dataset.

**Supplementary Figure 3.**
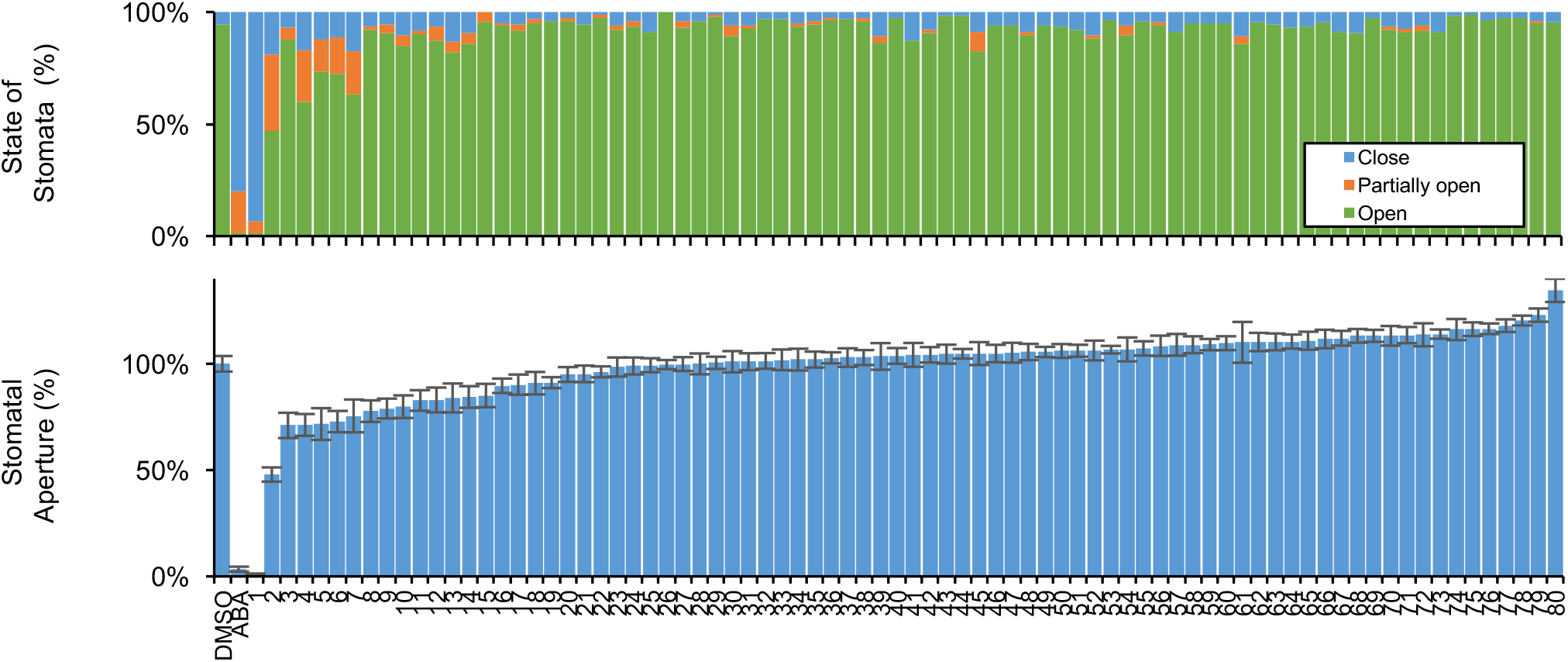
Summary of chemical screen to identify compounds that inhibit light-induced stomatal opening. Annotation follows that of Fig. 3d and 3e.

## References

1. Shimazaki, K., Doi, M., Assmann, S. M. & Kinoshita, T. Light Regulation of Stomatal Movement. Annu. Rev. Plant Biol. 58, 219–247 (2007).

2. Negin, B. & Moshelion, M. The evolution of the role of ABA in the regulation of water-use efficiency: From biochemical mechanisms to stomatal conductance. Plant Sci. 251, 82–89 (2016).

3. Zeng, W., Melotto, M. & He, S. Y. Plant stomata: A checkpoint of host immunity and pathogen virulence. Curr. Opin. Biotechnol. 21, 599–603 (2010).

4. Kutsuna, N. et al. Active learning framework with iterative clustering for bioimage classification. Nat. Commun. 3, 1032 (2012).

5. Tomiyama, M. et al. Mg-chelatase I subunit 1 and Mg-protoporphyrin IX methyltransferase affect the stomatal aperture in Arabidopsis thaliana. J. Plant Res. 127, 553–563 (2014).

6. Tsuzuki, T. et al. Mg-chelatase H subunit affects ABA signaling in stomatal guard cells, but is not an ABA receptor in Arabidopsis thaliana. J. Plant Res. 124, 527–38 (2011).

7. Hayashi, M., Inoue, S.-I., Takahashi, K. & Kinoshita, T. Immunohistochemical detection of blue light-induced phosphorylation of the plasma membrane H+-ATPase in stomatal guard cells. Plant Cell Physiol. 52, 1238–48 (2011).

8. Hayashi, M., Inoue, S., Ueno, Y. & Kinoshita, T. A Raf-like protein kinase BHP mediates blue light-dependent stomatal opening. Sci. Rep. 7, 45586 (2017).

9. Toh, S. et al. Identification and Characterization of Compounds that Affect Stomatal Movements. Plant Cell Physiol. (2018). doi:10.1093/pcp/pcy061

10. Amos, B., Ludwiczuk, B. & Satyanarayanan, M. OpenFace: A general-purpose face recognition library with mobile applications. C. Sch. Comput. Sci. Tech. Rep. C. 16, (2016).

11. Taigman, Y., Yang, M., Ranzato, M. & Wolf, L. DeepFace: Closing the gap to human-level performance in face verification. Proc. IEEE Comput. Soc. Conf. Comput. Vis. Pattern Recognit. 1701–1708 (2014). doi:10.1109/CVPR.2014.220

12. Cireşan, D. C., Meier, U., Masci, J., Gambardella, L. M. & Schmidhuber, J. ImageNet Classification with Deep Convolutional Neural Networks. Lancet (London, England) 346, 1501 (2011).

13. King, D. E. Max-Margin Object Detection. (2015).

14. Dalal, N. & Triggs, B. Histograms of Oriented Gradients for Human Detection. in 2005 IEEE Computer Society Conference on Computer Vision and Pattern Recognition (CVPR’05) 1, 886–893 (IEEE, 1994).

15. Schroeder, J. I., Allen, G. J., Hugouvieux, V., Kwak, J. M. & Waner, D. GUARD CELL SIGNAL TRANSDUCTION. Annu. Rev. Plant Physiol. Plant Mol. Biol. 52, 627–658 (2001).

16. Grys, B. T. et al. Machine learning and computer vision approaches for phenotypic profiling. 1–7 (2016). doi:10.1083/jcb.201610026

17. Girshick, R., Donahue, J., Darrell, T. & Malik, J. Rich feature hierarchies for accurate object detection and semantic segmentation. (2013).

18. Girshick, R. Fast R-CNN. (2015).

19. Ren, S., He, K., Girshick, R. & Sun, J. Faster R-CNN: Towards Real-Time Object Detection with Region Proposal Networks. (2015).

20. Liu, W. et al. SSD: Single Shot MultiBox Detector. (2015). doi:10.1007/978-3-319-46448-0_2

21. Long, J., Shelhamer, E. & Darrell, T. Fully Convolutional Networks for Semantic Segmentation. (2014).

22. Chen, L.-C., Papandreou, G., Kokkinos, I., Murphy, K. & Yuille, A. L. DeepLab: Semantic Image Segmentation with Deep Convolutional Nets, Atrous Convolution, and Fully Connected CRFs. (2016).

